# Intrinsic Repair Capacity of Resident Tendon Cells is Dependent on Hole Size in an Ex Vivo Model of Laser-Induced Microdamage

**DOI:** 10.1101/2025.05.29.656650

**Authors:** Anthony N. Aggouras, Matthew T. Lim, Jeroen Eyckmans, Brianne K. Connizzo

## Abstract

While it is generally accepted that tendon healing following widespread extracellular matrix trauma is limited, tenocytes are thought to have the capacity to repair small amounts of microdamage generated through activities of daily living. Despite this, few studies have directly studied the mechanisms governing this process. To address this, we developed a tunable *in vitro* model of extracellular matrix microdamage in live tendon explants that enables us to track both clearance of denatured collagen microdamage and closure of a micro-sized defect in the tendon matrix. The purpose of this study was to controllably induce varying levels of localized microdamage to the tendon explants and identify (1) if thresholds for healing exist and (2) whether repair mechanisms are dependent on initial damage size. We found that within three weeks, all tendon explants were able to clear damaged matrix to some extent regardless of the damage size. Interestingly, larger 5 mJ and 10 mJ injuries resulted in a more robust rate of damaged matrix clearance in the later weeks, while smaller injuries exhibited a more consistent rate that led to full clearance in two explants. Greater than 50% clearance of denatured collagen microdamage was typically associated with an accompanying closure of the ECM defect, suggesting a strong relationship between clearance and closure. Overall, our work demonstrates the power of our laser-induced microdamage model, which enables the direct visualization of microdamage responses. This model will be a powerful asset for investigating mechanisms of damage accumulation and/or healing, as well as identifying local tendon-specific factors that can be leveraged for therapeutics.

## Introduction

Tendinopathy is a chronic, degenerative condition marked by persistent pain, impaired function, and poor quality of life.^1,2^ It is thought to arise from a compromised ability to maintain and repair the extracellular matrix (ECM), resulting in the gradual accumulation of matrix damage that leads to macroscopic tendon tears.^3,4^ The structural, mechanical, and molecular changes associated with tendinopathy have been extensively characterized using fatigue loading and overload models.^5–9^ These studies highlight the susceptibility of tendon tissue to mechanical damage, as a single overloading event stretching the tendon past its yield point can induce ECM microdamage in the form of collagen kinks and denatured collagen.^9,10^ Despite the link between impaired ECM repair and tendinopathy, the underlying cellular mechanisms that support microdamage repair remain poorly understood.

Under physiological conditions, resident tendon cells maintain tissue structure and function through a delicate balance of matrix synthesis and degradation. In response to increased mechanical loading, such as during exercise, tenocytes actively engage in ECM remodeling by removing damaged matrix via upregulation of matrix metalloproteinases (MMPs), a disintegrin and metalloproteinase with thrombospondin motifs (ADAMTS), and cathepsins, followed by deposition of new matrix proteins.^11–13^ These neomatrix proteins are then incorporated and remodeled into the existing hierarchical structure to restore or improve mechanical function.^14^ While remodeling is well studied in the context of global changes in mechanical stimulus, such as exercise, much less is known about how tenocytes respond to localized ECM microdamage.

Microdamage repair has mainly been studied following widespread damage induced by repetitive fatigue loading. The most informative model to date was developed by Fung et al., in which the patellar tendon is clamped and directly loaded *in vivo*.^5^ Using this model, studies have suggested there is a limited capacity for tendon microdamage repair. Moderate levels of fatigue loading led to an immediate 20% stiffness loss and structural matrix damage that was unrecoverable even 8-10 weeks later.^15^ This was attributed to a lack of repair capacity in the resident tenocytes.^16^ In response to lower levels of fatigue-induced microdamage, tendons increased MMP and collagen expression at 7 days post-injury, suggesting that there was matrix turnover occurring.^17^ However, it’s not clear whether this constitutes microdamage repair or exercise-based adaptation. More recently, a few studies have looked at the innate repair response of tenocytes using *in vitro* models of induced microdamage.^18,19^ In these studies, tenocytes isolated from the systemic injury response appear to have a poor innate repair capacity to widespread matrix microdamage. While unable to recover the matrix damage, the tenocytes are still able to mount a robust repair response by releasing inflammatory markers and matrix degrading enzymes. However, the widespread damage used in these studies may be too extensive for an effective tenocyte-mediated local matrix remodeling response. Therefore, there is a need for a complementary model to study local tenocyte mediated ECM repair mechanisms in response to small levels of microdamage.

A promising method for inducing precise highly localized matrix damage in tendon is laser ablation, wherein a focused laser pulse directs a high amount of energy into a small area. Damage produced by laser ablation is often on the order of microns and the size of the damaged region can be tuned by altering the focusing angle and pulse energy of the laser.^20^ The ability to create tunable small micron-sized injuries with surrounding microdamage in the form of denatured ECM proteins makes laser ablation a powerful tool for studying cellular response to localized matrix damage. This technique has been used to observe how fibroblasts and other cell types embedded in engineered microtissues respond to ECM damage.^21,22^ By leveraging laser ablation to generate consistent microdamage, further insight was gained into the response of resident cells to injury and the mechanisms that mediate damaged ECM clearance. When used to recapitulate the acute ECM damage induced by a myocardial infarction, the cardiomyocytes and cardiac fibroblasts repair response mimicked an *in vivo* response characterized by cell death, granulation tissue formation, and early scar formation.^22^ Additionally, an increase in vimentin and fibronectin suggested that resident cardiac fibroblasts may be sufficient to drive some contraction of the tissue defect and deposition of new matrix. The laser ablation damage model was also employed to identify that the resident fibroblasts actively engulf and clear denatured ECM from the injury site, and this removal is dependent on ROCK and dynamin activity.^21^

To isolate and understand the local response of the resident tendon tissue and cells during matrix repair, tendon explants can be leveraged. Tendon explants allow for the study of extracellular matrix turnover within an isolated tissue while preserving cell–cell and cell–ECM interactions. Therefore, the objective of the present study was to establish a laser ablation technique to reliably induce various levels of microdamage “holes” in a live tendon explant, and then use this model to explore intrinsic repair independent from the influence of the systemic response. We hypothesized that resident tendon cells would be able to effectively repair and remodel microdamage on their own, and that resident repair capacity would be dependent on microdamage size.

## Methods

### Design of Laser Ablation Bioreactor

While laser ablation has been used to produce microdamage in microtissue constructs,^21,22^ it has not yet been used to produce damage to a whole tissue explant. This required the design and fabrication of a novel bioreactor to ablate and culture tendon explants while maintaining physiologic loading conditions. Our custom system was designed with a glass window in the bottom of the culture wells to allow for visualization of the explants during ablation (Figure 1A). The windows also include etched lines as a method for measuring the position of the ablated “hole” in reference to the grip edge (Figure 1A lower left inset). Tracking the position of the laser ablation site ensured the correct area of the tendon was analyzed, even if the damage induced by the laser was fully repaired. Inclusion of an in-line load cell and a linear variable differential transformer (LVDT) in the system also enables precise and controlled cyclic loading protocols on the explants, as previously described.^23^ The novel design is also tailored for the ablation scope and allows for visualization, ablation, and culture of up to 2 tendon explants in each well (Figure 1A). The ability to ablate the explants directly in the bioreactor avoids added handling and reduces the amount of time that tendons remain outside of culture. The bioreactor has 6 center wells which allow for ablation (12 tendons) and 4 outer wells which can serve as wells for control non-ablated tendons (8 tendons).

**Figure 1.**
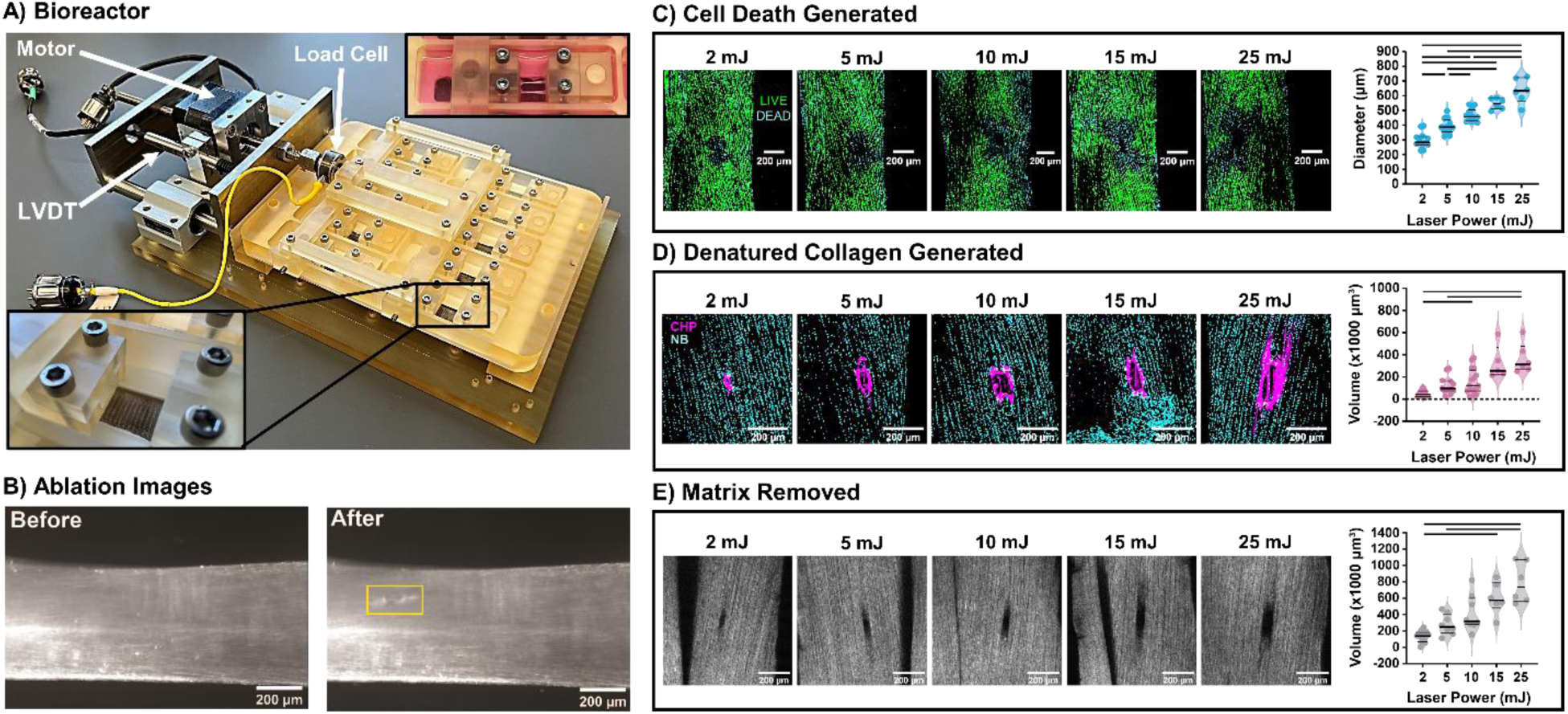
Characterization of the ablation system. (A) An image of the laser ablation bioreactor with the hardware for measuring and controlling strain and load on the explant. Explants are loaded two per well into the grips (inset, upper right), and the site of ablation is tracked relative to grip position through etched lines on the glass bottom wells (inset, lower left). (B) Before and after images of an ablated tendon with a yellow box highlighting the area of damage after single 10 mJ pulse ablation. (C) Cell nuclei stained with fluorescein diacetate (live cells only, green), and NucBlue (all cells, cyan) demonstrate increasing cell death with increasing laser power (D) Denatured collagen stained via CHP (magenta) increases with increasing laser power (E) Increase in ECM removal with increasing laser power, where removal of collagen ECM presents as a dark hole in the SHG signal. All data is presented with median and quartiles shown as black lines on violin plots with individual data points. Significant comparisons between laser powers are denoted as a bar (-) p < 0.05.

### Sample Preparation

This study utilized the flexor digitorum longus (FDL) tendon harvested from the limbs of skeletally mature young (2-4 months) C57BL/6J male mice directly following sacrifice per approved animal use protocol (BU IACUC PROTO202000046) as the explant model. Following previously described methods,^24^ all explants were washed in 1X PBS supplemented with antibiotics (100 units/mL penicillin G, 100 µg/mL streptomycin (Fisher Scientific, Waltham, MA), and 0.25 µg/mL Amphotericin B (Sigma-Aldrich)). Explants were then immediately loaded into bioreactor grips using a loading rig to ensure all tendons were reliably held at a 10-mm gauge length. The tendons were oriented such that the joint-facing side of the tendon was ablated. The gripped tendons were then placed into the custom-built tendon laser ablation bioreactor and stretched manually to approximately 1% strain, which has been shown previously to be adequate for maintenance of tissue properties.^23^ Left and right tendons from every mouse were used to conserve animal numbers, but left and right tendons from the same animal were not attributed to the same experimental group. Throughout ablation and culture, explants were maintained in standard culture medium consisting of low glucose Dulbecco’s Modified Eagle’s Media (1 g/L; Fisher Scientific) supplemented with 10% fetal bovine serum (Cytiva, Marlborough, MA) and antibiotics.

### Ablation

For ablation, the bioreactor was transported to the stage of a Zeiss Axiovert S100 microscope equipped with a 10x objective and a 1064 nm q-switched pulsed-nanosecond Nd:YAG laser (Minilite, Continuum, San Jose, California). As previously described,^21^ the beam is enlarged by a beam expander (Edmund Optics #39-739, Barrington, New Jersey) from 3 to 9 mm, directed through a dichroic mirror suitable for 1064 nm, and focused into the tissue through a 10x objective. For this study, laser energy was measured directly after the beam expander and was adjusted to the desired laser power: 2, 5, 10, 15, or 25 mJ. All tendons were ablated near the bifurcation of the two fascicles of the FDL, in the middle of the larger fascicle. To accomplish uniform ablation, the microscope was focused on the outer edge of the large fascicle to ensure a uniform depth of the laser focal point. Phase images were taken at 10x with an Axiovert inverted microscope (Zeiss, Oberkochen, Germany) equipped with a 20Mp Blackfly camera (FLIR, Wilsonville, Oregon) before and immediately following injury. Using the Zeiss Axiovert S100 microscope, tendons were visualized before and after microdamage (Figure 1C). The ablated microdamage “hole” shown in the representative brightfield image was produced with a single 10mJ laser pulse (Figure 1B).

A non-ablated control injury (‘Puncture’) was also developed to account for differences in repair capacity attributed to heat damage generated by the laser. A 32-gauge needle was pushed through the large fascicle of the explant at a similar location along the length of the tendon as the ablations. The 32-gauge puncture produces a “hole” defect comparable to the 5mJ laser pulse without producing heat-denatured collagen in the process.

### Cell Viability

Cell viability was assessed through live/dead staining in 1X PBS containing fluorescein diacetate (FDA; 4 mg/mL; Fisher Scientific) and NucBlue nuclear stain (2 drops/mL; Thermo Fisher Scientific, Waltham, Massachusetts). Dead cells were identified as positive for NucBlue staining but negative for FDA staining. To obtain the cell death area following ablation, a circular ROI was drawn containing all dead cells around the injury. The diameter of that circular ROI was used to compare the extent of cell death generated by the laser ablation.

### Quantification of Damaged ECM and Clearance

During ablation, the highly localized energy is dispersed as heat and creates thermal denaturation of surrounding ECM proteins. Accompanying the thermal denaturation is a local pressure buildup in the tissue that results in mechanical failure of the surrounding ECM and destructive tissue ablation,^20^ leaving an ECM defect in the form of a “hole”. The denaturation of collagen by the thermal energy released in the tissue during ablation allows the damage generated to be stained via collagen hybridizing peptide (CHP). Therefore, collagen damage and cell location were determined by Cy3-CHP (4.32 μg/mL; Advanced Biomatrix, Carlsbad, California) with NucBlue (2 drops/mL; Thermo Fisher Scientific) counterstain, respectively.

Briefly, a 200 μL CHP solution of 4.32 μg/mL CHP was heat shocked at 80 °C for 10 minutes and then cooled in a water bath for 1.5 minutes. After cooling, tendons were added to the solution and stained at 4 °C for 18 hours. Tendons were then removed from the CHP stain and transferred to a NucBlue stain containing 2 drops of NucBlue stock in 1 mL of PBS for 5 minutes. Confocal image z-stacks of the ablation site were captured on an Olympus FV3000 with a 20x objective.

Custom software was used to quantify damage intensity and dimensions, as well as the number and area of cell nuclei in the damaged region. First, the background signal was removed through user-defined thresholding of the stack of CHP images. The max projection image of the image stack was then used to identify the region of interest (ROI). The ROI was defined by the code as the pixels that fell within CHP stain regions verified by user input. The depth of the denatured collagen was assessed by identifying the number of slices containing CHP stain in the ROI. The width and the length of the denatured collagen ROI were obtained by taking the distance between ROI edges in the direction perpendicular and parallel to the long axis of the tendon, respectively. The volume of denatured collagen was determined by quantifying the area of CHP stain in the drawn ROI on each plane of the image stack and multiplying by the image slice height of 3.94 μm. Average collagen damage intensity was determined by summing the intensity of each pixel of the ROI across all slices, including pixels without stain, and taking the average of the summed intensity of all the pixels in the ROI.

For studies tracking changes in matrix microdamage over time, clearance is presented as the percent difference from initial injury (day 0) to account for differences in initial damage size between laser powers. The daily rate of clearance is presented as the change from the previous time point divided by seven days. These measures are shown for denatured collagen depth, width, length, and average intensity in the supplement.

### Quantification of Hole Morphology and Closure

The mechanical failure of the surrounding ECM and destructive tissue ablation leaves a “hole” in the tendon ECM that is detectable via Second Harmonic Generation (SHG) imaging. Tendons were imaged for SHG using a two-photon microscope with the laser tuned to 1190 nm, a dwell time of 1.2 ms, and an imaging window of 1024 x 1024 pixels. Pockels were interpolated exponentially from 130 at the tendon surface to 180 at 150 µm into the tendon, with frame-averaging every two frames. Custom software was then used to quantify the volume of the “hole” left in the tendon ECM. Briefly, a gaussian filter was applied to the SHG images to reduce noise, and the images were auto-thresholded using a locally adaptive threshold with high sensitivity. This ensured low level signal was kept, but noise was removed. The user then selected the frame where the ECM “hole” first begins, the frame where clear collagen fibers disappear, and the position of the ECM “hole”. The software is then able to track the hole across the z-stack and calculate the number of pixels without SHG signal. “Hole” volume is then calculated by quantifying the area of continuous dark pixels on each plane of the image stack and multiplying by the image slice height.

For studies tracking changes in hole size over time, closure is presented as the percent difference from initial injury (day 0) to account for differences in initial hole size between laser powers.

### MMP Activity

To understand how the resident cells were facilitating the removal of denatured collagen from the injury site, we assessed matrix metalloproteinase (MMP) activity. MMP (1,2,3,7,8,9,10,13,14) activity was determined via analysis of spent culture medium (n= 2- 6/group/time point) using a commercially available FRET-based generic MMP cleavage kit (SensoLyte 520 Generic MMP Activity Kit Fluorimetric, Anaspec, Fremont, CA). Media was collected during standard media changes on days 6, 14, and 20 of culture. MMP activity is represented as the concentration of MMP cleaved product (5-FAM-Pro-Leu-OH), the final product of the MMP enzymatic reaction.

## Data and Statistics

All data are presented as violin plots with median, as well as first and third quartiles marked. Data points more than 2 standard deviations outside of the mean were removed as outliers. Statistical evaluation on this set of data was performed in GraphPad Prism 8 (GraphPad, San Diego, CA) using individual one-way ANOVAs for each laser power level with effects for time in culture. Bonferroni corrected post-hoc t-tests were then used to identify differences within each time point where appropriate. Bartlett’s test for heteroscedasticity and D’Agostino-Pearson omnibus test for normality were used to assess the data sets. In instances where a data set was heteroscedastic, a Brown-Forsythe ANOVA and Dunnett’s T3 multiple comparisons test were used for statistical evaluation. In instances where a data set was not normally distributed, a Kruskal-Wallis test and Dunn’s multiple comparisons test were used for statistical evaluation. A significant increase in the rate of clearance was determined through a one-tailed t-test of the timepoint compared to a hypothesized zero rate of clearance. For all comparisons, significance was noted at *p<0.05. Correlations between the denatured ECM cleared and the ECM “hole” closure were assessed using simple linear regression analysis.

## Results & Discussion

To identify the capacity of resident tendon cells to initiate local matrix repair of various levels of microdamage, methods were established to create tunable local ECM microdamage in the tendon explants. As previously shown in other biological tissues,^20^ adjusting the laser pulse energy enabled multiple sizes of microdamage to be produced with varying affected damage dimensions. As laser pulse energy is increased, the diameter of the region of cell death also increases steadily (Figure 1C). Denatured collagen volume increased with an increase in laser power (Figure 1D), most likely a result of significant increases in damage width (transverse dimension of the tendon) and damage depth (Supplemental Figure S1). Damage length (axial dimension of the tendon) remained largely unchanged except at high laser powers where it appeared to increase (not statistically significant, Supplemental Figure S1). Accompanying this increase in the volume of denatured collagen was an increase in the volume of the ECM defect, which is characterized by a detectable “hole” in a SHG image of the ablation site (Figure 1E). Together, this demonstrates successful creation of a <u>tunable</u> and <u>repeatable</u> micron-sized damage region with mechanical failure of the collagen matrix and heat denaturation of matrix proteins in a live tendon explant. The ability to scale the affected area by adjusting laser pulse energy and the potential inclusion of multiple damage sites allow for a customizable approach for studying resident cell responses to localized microdamage.

The next goal was to explore how initial injury size affects intrinsic repair capacity. To create three distinct microdamage sizes, tendon explants were ablated with a laser power of 2, 5, or 10 mJ. Representative images show a progressive reduction in CHP stain around the ablated hole during the culture period for all damage levels (Figure 2A), suggesting that resident cells can remove damaged ECM without extrinsic support.

**Figure 2.**
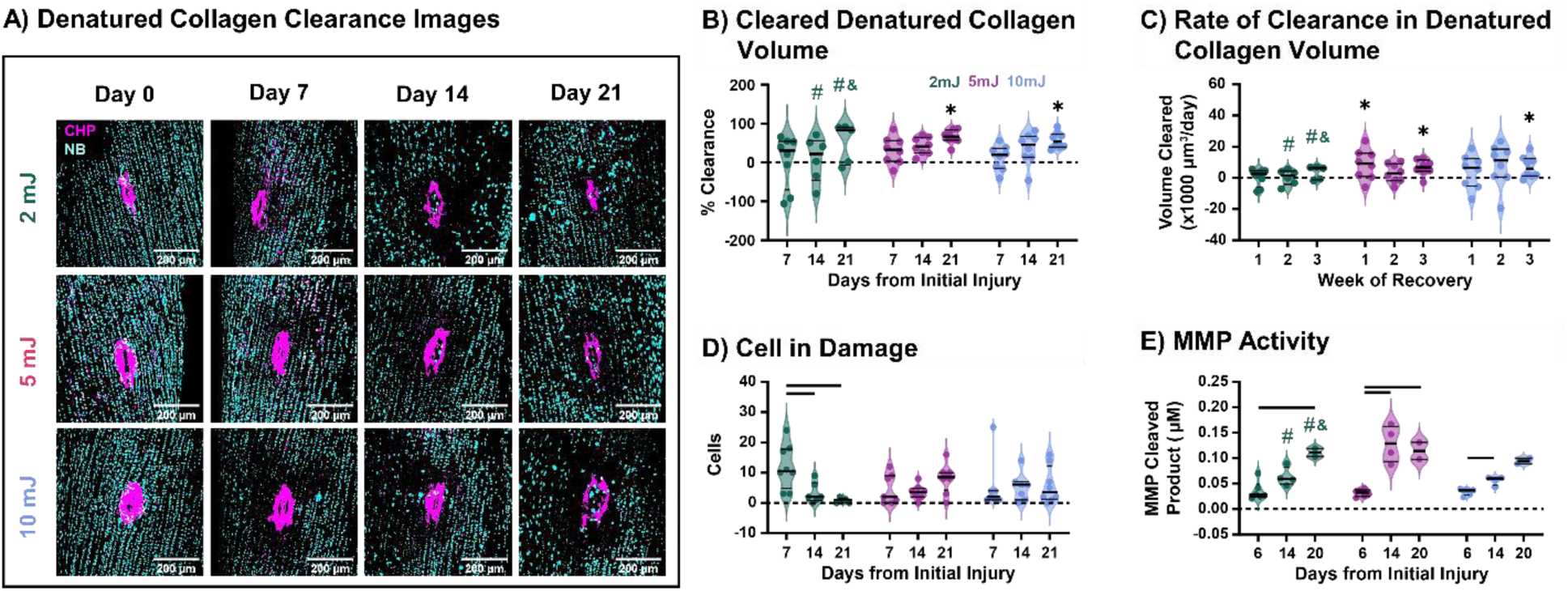
(A) Representative max intensity projection images of collagen damage (magenta) and cell nuclei (cyan). Damage from 2 mJ (top row), 5 mJ (middle row), and 10 mJ (bottom row) at days 0, 7, 14, and 21. (B) Percentage of total clearance and (C) weekly rate of clearance in denatured collagen damage volume. (D) Cells present within the bounds of the denatured collagen damage. (E) Generic MMP activity of explants measured from culture media taken at media changes close to takedown time points. In the 2mJ group one tendon on day 14 and one on day 21 had fully cleared during culture (represented by #). In the 2mJ group, two tendons on day 21 had degraded during culture (represented by &). Data is presented with median and quartiles shown as black lines on violin plots with individual data points. Significant comparisons (p < 0.05) between timepoints are denoted as a bar (-) spanning between time points, and comparisons to baseline (day 0) are denoted by a star (*).

Using the developed custom software, the clearance of denatured collagen was measured by the reduction in CHP volume over the culture period. The amount of cleared denatured collagen measured by CHP stain volume increased over time in culture for all groups but was significant from the baseline at day 21 in the 5 mJ and 10 mJ groups only (Figure 2B). Of note, two tendons subjected to 2 mJ injury were able to fully clear the denatured collagen over the three-week culture period, with one clearing by day 14 of culture and the other by day 21. Two tendons in the same group had also degraded by day 21, marked by “frayed” fibers and indicating potential excessive removal of the collagen matrix beyond just damaged ECM. Only the 5 and 10 mJ groups exhibited an increased rate of clearance over one or more weeks in culture (Figure 2C). The 2 mJ group had a consistent low rate of clearance across all weeks, while the 5 mJ group had an increased rate of clearance over the first and third week, and the 10 mJ group only had an increased rate of clearance over the third week. Together, this suggests the timing of denatured collagen clearance may vary based on the size of injury, but the extent and ability to clear denatured collagen remains consistent across injury size.

All groups exhibited a low level of cellularity at the injury site (Figure 2D). In the 2 mJ group, there was a decrease in the number of cells at the injury site from day 7 to days 14 and 21. The early presence of cells at the injury site in the 2 mJ group may have contributed to the full clearance of two tendons by day 14 and 21. In contrast, the 5 and 10 mJ groups exhibit slight increases over culture. The 5 mJ group also appears to have some explants that have around 10 cells in the damage at day 7, which could explain why there is a heightened rate of clearance in the 5 mJ group over the first week of culture. The 10 mJ group had more cells present in the damage on days 14 and 21, which could account for the increased clearance seen in some explants over the last two weeks of culture. Cell death and matrix damage induced by the larger injuries could have slowed cell migration, resulting in the differences seen in cell infiltration to the injury site. These differences in the presence of cells across the groups may account for the influence that initial injury size has on the timing of denatured collagen clearance. Nevertheless, the conserved presence of cells within the bounds of the laser ablation induced microdamage at all damage levels suggests that the clearance of acute microdamage is an intrinsic cell-mediated process. Studies on non-viable tendons did not demonstrate any clearance of the denatured collagen over 21 days in culture, confirming that live cells are necessary for this process (data not shown).

While the cells present at the damage site have not been identified, they may be cells that migrate from other areas of the tendon explant due to inflammatory signaling following the injury. In acute injuries, resident tendons cells can initiate a pro-inflammatory environment and the recruitment of immune cells within 2-3 days through the release of cytokines such as interleukin-1β, tumor necrosis factor-α, and interleukin-6.^25^ Future studies plan to quantify the levels of secreted pro-inflammatory cytokines near the injury site. Regardless, it’s possible these cells belong to a resident immune cell population, such as resident macrophage-like cells.^26^ Given the small population of these cells, it seems equally likely that they are migratory tenocytes responding to inflammatory signaling.^27^ Regardless of their origin, which current studies are exploring, it’s clear that cells near the microdamage site play an important role in the clearance of denatured collagen microdamage.

We then questioned how cells at the injury site are coordinating this robust clearance of damaged tissue. The removal of damaged matrix has been associated with the expression of MMPs, ADAMTs, and cathepsins in other injury models.^12,13^ However, we found that MMP activity increased over the culture period (Figure 2E) in all groups, coinciding with an increase in the percentage of cleared denatured collagen but not with the rate of clearance of denatured collagen. However, our assay is non-specific, suggesting it’s possible that only specific MMPs are contributing to the clearance of denatured collagen such as MMP-1,-2,-8,-9 and-13.^12,28^ This method also does not look at local presence of MMPs but the activity of secreted MMPs from the whole explant, therefore local expression of MMPs may show a better correlation to the rate of denatured collagen clearance. Future studies will seek to identify the local presence of specific proteases, including MMPs, ADAMTs, and cathepsins to identify how the cells are driving denatured collagen clearance. Furthermore, there could be other processes that have not yet been explored. Previous studies have shown that fibroblasts have the capacity to recycle collagen through an endocytic process and that removal of denatured ECM from an injury site relies on ROCK and dynamin activity, suggesting a phagocytic or endocytic process.^21,29^

Interestingly, there was a significant increase in MMP activity from day 6 to 14 in the 5 mJ and 10 mJ groups, which was not present in the 2 mJ injury. Tendon is a mechanosensitive tissue, and the ability of the tendon cells to respond to a loss in mechanical cues is an important part of the healing cascade.^30,31^ It is important to note that the expression of MMPs responsible for clearing denatured collagen from the damage area is known to be affected by mechanical loading.^18,31–33^ The larger injuries produced by the larger laser pulses may lead to more unloading of collagen fibrils and fibers in the tissue due to disrupted matrix, which is known to increase MMP production in tendon.^24,32,34,35^ Future studies will investigate the local mechanical environment near the ablation site to ascertain if the local tendon mechanics influence protease activity after microdamage injury.

Following clearance of damaged collagen, cells then facilitate the deposition and replacement of missing collagen matrix. The initial “hole” left in the collagen matrix by the individual laser pulses and puncture injury, as well as any potential resident tendon cell mediated repair or infill of the “hole” after 21 days of culture, was visualized through SHG imaging. (Figure 3A). In response to an injury in which the tendon ECM is removed, resident tendon cells will produce neomatrix such as collagen III and collagen I,^36^ both of which will show up on tendon SHG imaging.^37^ As stated previously, the 32-gauge puncture injury served as a reference for ECM “hole” defect closure in an injury model that did not generate heat-denatured collagen, with comparable hole volume to a 5 mJ laser pulse. The puncture group closed the ECM hole to a similar extent as the laser injury groups, indicating that the presence of denatured collagen does not affect ECM “hole” repair.

**Figure 3.**
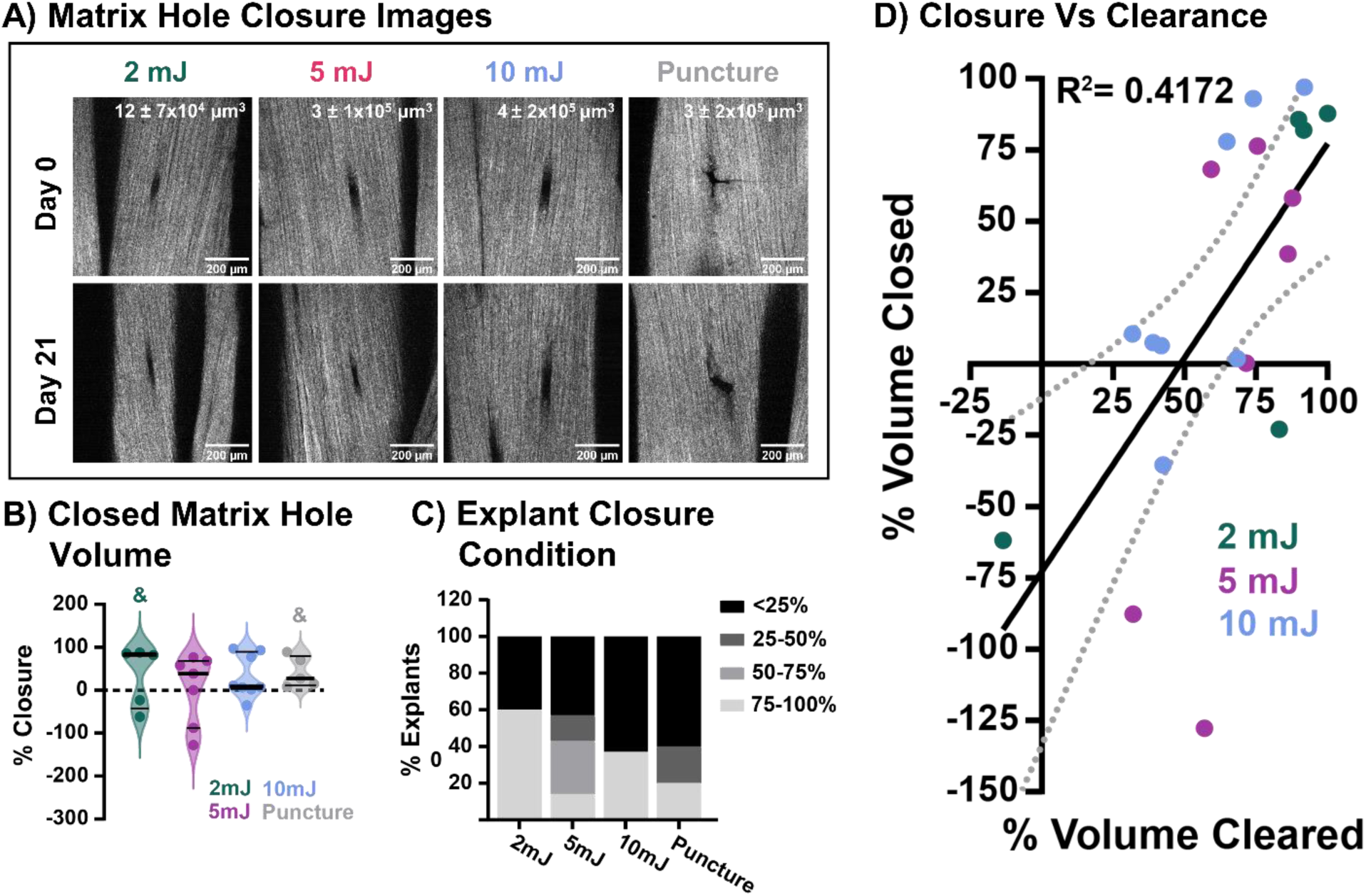
(A) SHG representative images of collagen ECM removed by laser ablation 2 mJ (left column), 5 mJ (middle left column), 10 mJ (middle right column), and by 32-gauge needle puncture (right column) at days 0 and 21. Numbers in top right of day 0 images show initial defect volume. (B) Percentage of total ECM hole closure over the 21 days of culture. Two tendons in the 2mJ group and one tendon in the puncture group had degraded by day 21 of culture (represented by &). Data is presented with median and quartiles shown as black lines on violin plots with individual data points. (C) Percentage of explants in each quartile of closure. (D) Paired closure of ECM hole versus clearance of denatured collagen from the injury site for each sample represented as a percentage. Linear regression with 95% confidence intervals represented by black line. R-squared value of the linear regression shown on the plot.

No injury group showed significant percent closure of the collagen matrix “hole” from their respective initial “hole” volume (Figure 3B). However, categorizing the tendon explants by percent closure, it becomes clear that as initial injury size increases, fewer explants can initiate closure, with a greater percentage of explants staying below 50% closure (Figure 3C). The reduced closure in larger injuries could be due to the increased cell death seen in the larger injuries. Therefore, with more viable cells near the injury site, the explants with smaller injuries may have been able to start laying down neomatrix and contract existing matrix sooner, accounting for the increased number of tendons with a greater extent of detectable closure via SHG imaging. Alternatively, the closure observed may be due to a contraction of the surrounding matrix in response to laser ablation damage induced changes in the local matrix mechanical environment. Resident tendon cells have been shown to be able to drastically contract the ECM in response to changes in mechanical load through an α-smooth muscle actin-mediated cellular mechanism.^38^ While we believe that closure is a result of neomatrix deposition, we are exploring all options currently. Future studies will aim to identify the presence of contractile cells and the deposition of neomatrix proteins like collagen III and fibronectin.

Notably, some explants in the 2 mJ and 10 mJ group either had nearly complete closure (above 75% reduction of the initial “hole” volume) while others had not closed (below 25% reduction), and in some cases had increased above the initial “hole” volume. This could indicate a “fate switch” for hole closure, resulting in this bimodal distribution. Several factors could contribute to this phenomenon, including initial cell death, mechanical environment, aberrant or insufficient neomatrix production, or insufficient cell types present during repair. In addition, the extent to which the resident tendon cells can clear damaged matrix may directly influence hole closure. Looking at the relationship between damage clearance and hole closure, it is clear that to have closure over 75%, there must also be at least 50% clearance of denatured collagen (Figure 3D). In fact, the amount of defect closure in the explant is moderately correlated with the amount of denatured collagen (R^2^= 0.4172). This would suggest that before the defect can be contracted or new collagen matrix can be deposited at the injury site, the injury site must be debrided of damaged and denatured ECM. We are exploring models of superior and inferior healing to identify key factors in both clearance and closure, and the relationship between these two processes. Regardless, this study has shown that across different extents of localized tendon ECM microdamage, resident tendon cells have the capacity to initiate repair through clearance of denatured collagen and, in some cases, are able to fully repair the collagen matrix with closure of the ECM defect.

This study and model system are not without their limitations. Injuries to tendons that generate microdamage are typically mechanical injuries, such as acute overload or overuse fatigue injuries. While heat-induced denaturation is not physiologic, this model is much more controllable and allows for easy quantification of initial damage and clearance over time. We have also determined that the microdamage generated by the laser ablation is on the scale of the microdamage generated during overload and overuse. Assuming the tendon is an elliptical cylinder with a width of 1mm and a thickness of 300 µm, the damaged area for 25 mJ laser pulse is about 0.47% of the total surface area of the 10mm cultured portion of the tendon explant. In overuse fatigue and overload injuries, collagen microdamage can account for 0.2-2 % of the total collagen and as high as 13% of the total tendon area.^6,10^ The average size of “microtears” in tendon can range from 0.33 to 3% in loaded tendons to 0.44 to 2.66% in unloaded tendons.^39^ Therefore, our model is able to generate denatured collagen that is common in tendon injuries at levels at the lower end of typical injury levels. Another limitation of the study is that the tendons are maintained at 1% static strain, rather than more physiologically relevant cyclic loading. We have found previously that a low-level static strain is sufficient to maintain the tendon during culture,^23^ but we also have ongoing studies to explore how loading influences the microdamage repair rate in the tendon explants following localized microdamage. Finally, this study is limited by the fact that all the assays used are endpoint assays. We were unable to track the same injury with the same geometry and cell environment across all time points, adding variability to our measurements. While we are confident that the initial injury conditions are highly consistent across the cultured explants, we are currently working to build a longitudinal system to enable real-time tracking of cell behavior, damage clearance, and neomatrix deposition. In addition, we are currently performing spatial transcriptomic and proteomic studies to better understand the local healing mechanisms and the important cellular pathways of resident tendon cell-mediated repair.

Despite these limitations, we present here a novel model for studying innate tissue healing processes following a microscale injury without interference from extrinsic factors. To our knowledge, our model system has, for the first time, provided the ability to directly identify that intrinsic cell-mediated responses may be sufficient to initiate ECM repair in response to localized microdamage injury. We have also identified that larger levels of microdamage affect the microenvironment enough to alter the timing of matrix repair, with delayed increases in the rate of damage clearance. We have also shown that closure of a microdamage “hole” defect may rely on the extent of damage clearance. Nevertheless, our novel laser ablation induced microdamage model enables the direct visualization and study of local tissue repair, which will be a powerful asset for understanding mechanisms of microdamage accumulation and/or healing, as well as identifying local tendon-specific factors that can be leveraged for therapeutics.

## Acknowledgements

This study was supported by NIH/NIA R00-AG063896, R21-EB028491, and MIRA: R35-GM151127. Research reported in this publication was supported by the Boston University Micro and Nano Imaging Facility and the National Institutes of Health (S10OD024993).

## Supplement

**Supplemental Figure S1.**
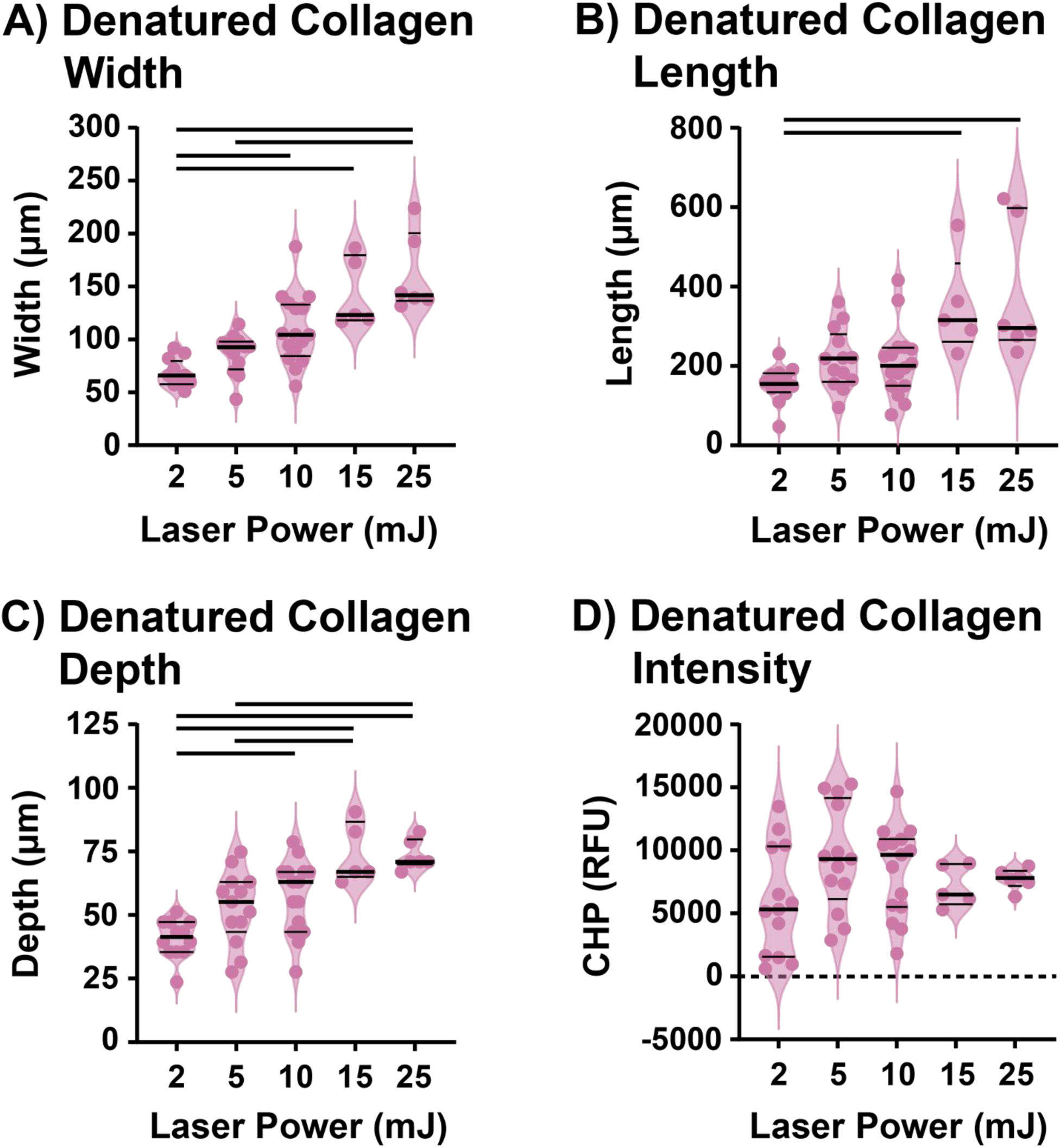
Analysis of initial denatured collagen in (A) width (transverse axis), (B) length (axial axis), (C) depth into the tendon, and (D) average pixel damage intensity when summed over all depths. All data is presented with median and quartiles shown as black lines on violin plots with individual data points. Significant comparisons between laser powers are denoted as a bar (-) p < 0.05.

**Supplemental Figure S2.**
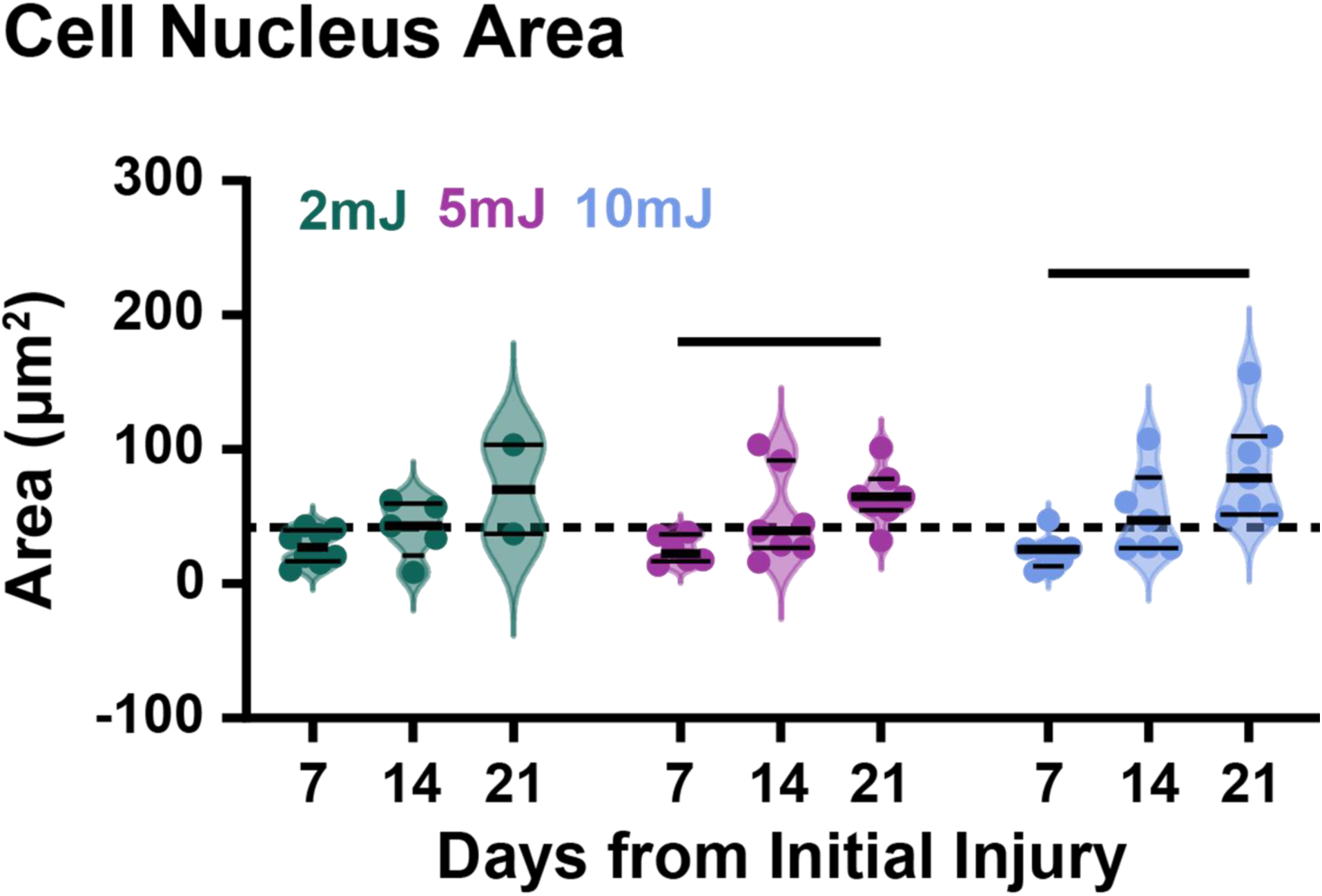
Cell nucleus area for 2 mJ (green), 5mJ (red), or 10 mJ (blue). Dashed line represents uninjured tenocytes mean nuclear area. Significant comparisons (p < 0.05) between timepoints are denoted as a bar (-)

**Supplemental Figure S3.**
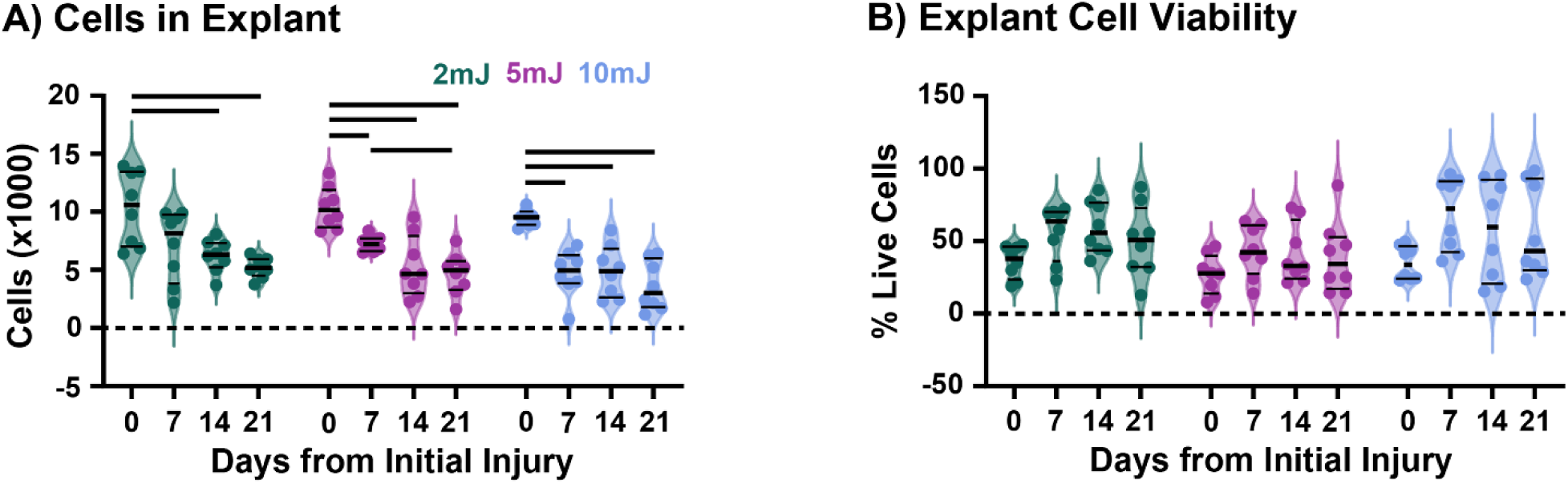
Analysis of cells present in the whole explant over culture. (A) Cell numbers and (B) explant viability represented as percent of alive cells in the explant for 2 mJ (green), 5mJ (red), or 10 mJ (blue) at days 0, 7, 14, and 21. Data is presented with median and quartiles shown as black lines on violin plots with individual data points. Significant comparisons (p < 0.05) between timepoints are denoted as a bar (-) spanning between time points.

**Supplemental Figure S4.**
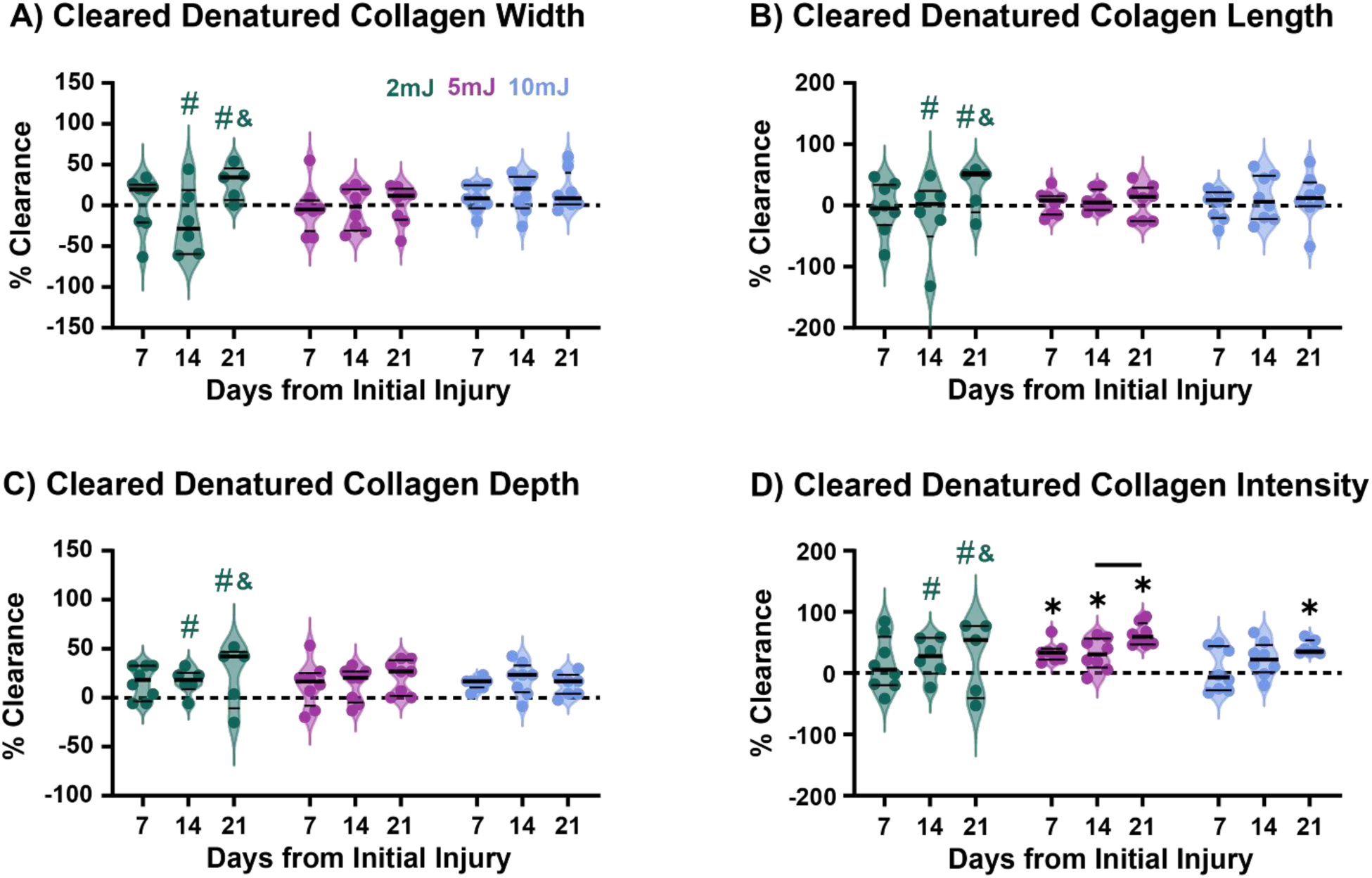
Percentage of total clearance of denatured collagen in (A) width (transverse axis), (B) length (axial axis), (C) depth into the tendon, and (D) average pixel damage intensity when summed over all depths. In the 2 mJ group one tendon on day 14 and one on day 21 had fully cleared during culture (represented by #). In the 2 mJ group two tendons on day 21 had degraded during culture (represented by &). Data is presented with median and quartiles shown as black lines on violin plots with individual data points. Significant comparisons (p < 0.05) between timepoints are denoted as a bar (-) spanning between time points, and comparisons to baseline (day 0) are denoted by a star (*).

**Supplemental Figure S5.**
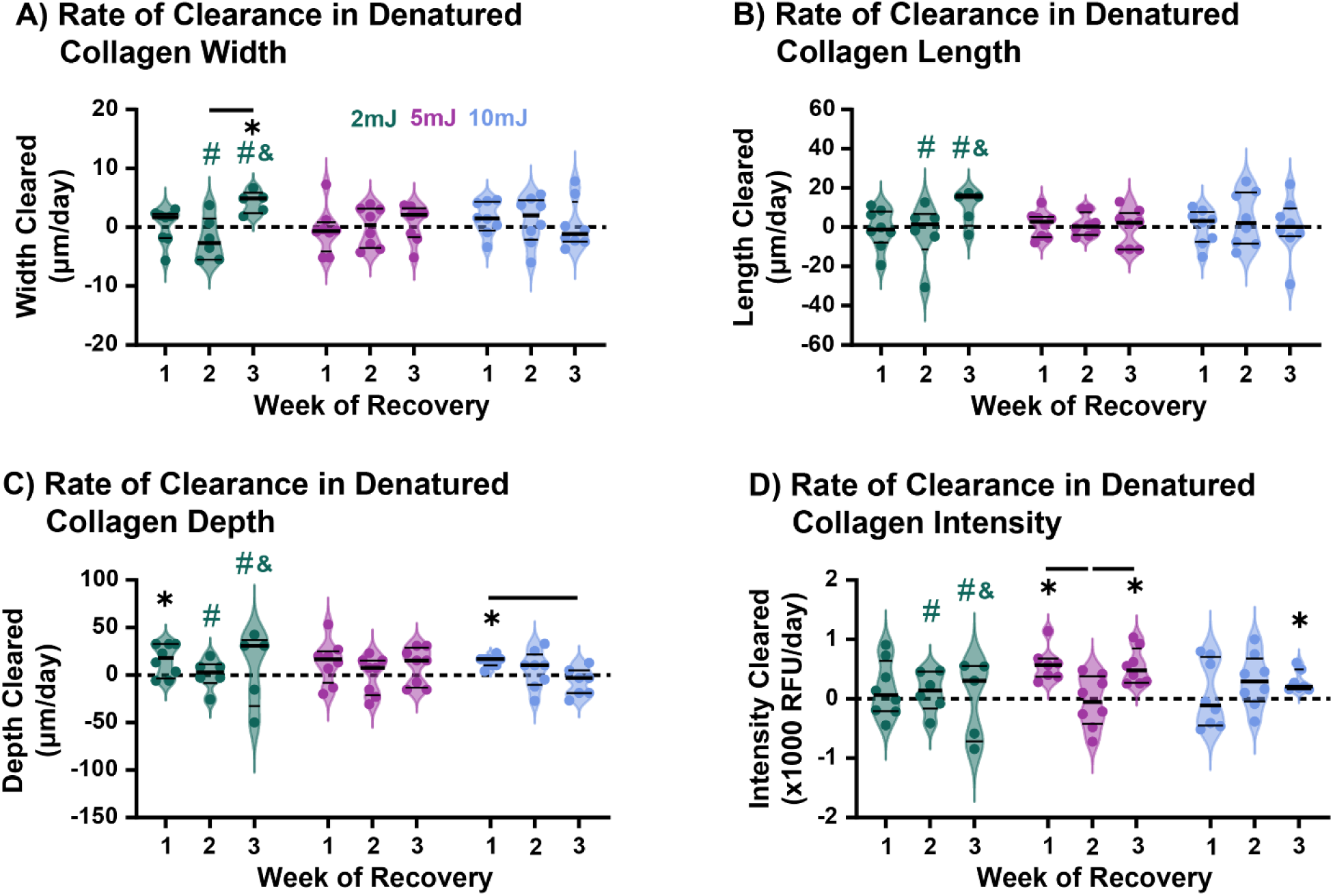
Weekly rate of denatured collagen clearance in (A) width (transverse axis), (B) length (axial axis), (C) depth into the tendon, and (D) average pixel damage intensity when summed over all depths. In the 2 mJ group one tendon on day 14 and one on day 21 had fully cleared during culture (represented by #). In the 2 mJ group two tendons on day 21 had degraded during culture (represented by &). Data is presented with median and quartiles shown as black lines on violin plots with individual data points. Significant comparisons (p < 0.05) between timepoints are denoted as a bar (-) spanning between time points, and comparisons to baseline (day 0) are denoted by a star (*).

**Supplemental Figure S6.**
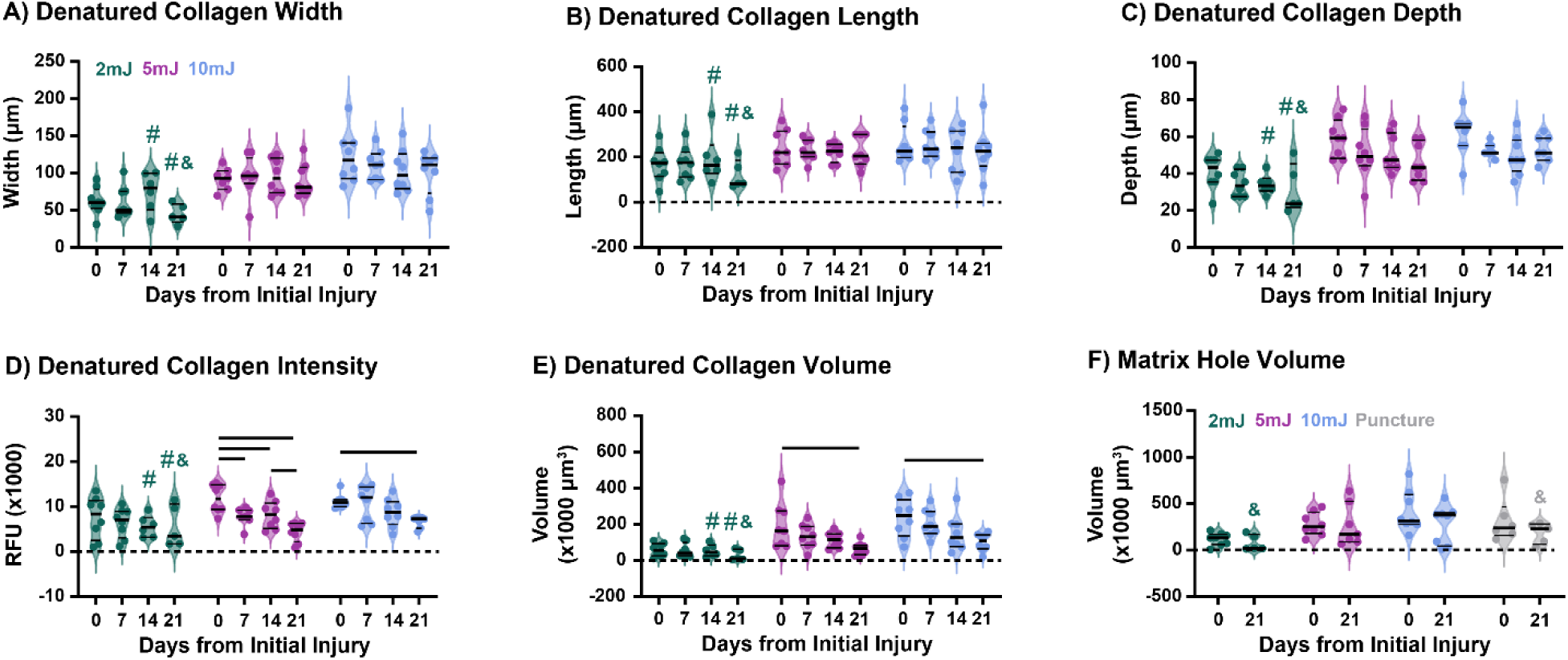
Amount of denatured collagen in (A) width (transverse axis), (B) length (axial axis), (C) depth into the tendon, (D) average pixel damage intensity when summed over all depths, and (E) volume at all time points. (F) Initial (Day 0) and final (Day 21) “hole” volume measured via SHG. In the 2 mJ group one tendon on day 14 and one on day 21 had fully cleared during culture (represented by #). In the 2 mJ group two tendons on day 21 had degraded during culture (represented by &). Data is presented with median and quartiles shown as black lines on violin plots with individual data points. Significant comparisons (p < 0.05) between timepoints are denoted as a bar (-) spanning between time points.

